# Genome-wide identification of stable RNA-chromatin interactions

**DOI:** 10.1101/2024.09.04.611281

**Authors:** Xingzhao Wen, Sheng Zhong

## Abstract

RNA-chromatin interactions play crucial roles in gene regulation and genome organization, but the interaction landscape remains poorly understood. In this study, we conducted an in-depth analysis of a previously published dataset on RNase-treated in situ mapping of the RNA–genome interactome in human embryonic stem cells. This dataset globally profiles RNase-insensitive RNA-chromatin interactions. Our analysis revealed that RNase treatment selectively preserved long-range RNA-chromatin interactions while removing promiscuous interactions resulting from the local diffusion of nascent transcripts. RNase-insensitive chromatin-associated RNAs (RI-caRNAs) exhibited high sequence conservation and preferentially localized to functional genomic regions, including promoters, transcription factor binding sites, and regions with specific histone modifications. Interestingly, coding and non-coding RNA transcripts showed distinct sensitivities to RNase, with lncRNAs and disease-associated transcripts being enriched among RI-caRNAs. Furthermore, we identified specific caRNA classes associated with individual transcription factors and histone modifications. Altogether, our findings reveal a RNase-inaccessible regulatory RNA-chromatin interactome and provide a resource for understanding RNA-mediated chromatin regulation.

## Introduction

The nucleus is a complex environment where RNA plays diverse roles beyond its canonical function in gene expression. These roles include the formation and maintenance of phase-separated condensates (Hnisz et al. 2017), modulation of 3D genome organization (Saldaña-Meyer et al. 2019; Hansen et al. 2019; Calandrelli et al. 2023) , facilitation of protein-chromatin interactions (Long et al. 2020; Sigova et al. 2015), and regulation of gene expression through enhancer or promoter upstream RNAs (T.-K. Kim et al. 2010; Liang et al. 2023). Many of these functions involve interactions between RNA, proteins, and specific regions on chromatin (Quinodoz et al. 2021; Xiao et al. 2019). Therefore, understanding the full spectrum of RNA functions in the nucleus requires elucidating both the interactions and localization patterns of nuclear RNAs.

Current methods for investigating nuclear RNA functions fall into two main categories. The first approach focuses on elucidating RNA function through its interactions with other molecules, such as RNA-protein interactions (Van Nostrand et al. 2016; Hafner et al. 2010) or RNA-RNA interactions (Lu et al. 2016; Nguyen et al. 2016). The second approach aims to understand RNA function through its localization pattern inside the nucleus. This is achieved either by probing the chromatin regions that RNAs are spatially close to using proximity ligation-based methods (Li et al. 2017; Sridhar et al. 2017; Wu et al. 2019) or ligation-free methods (Quinodoz et al. 2021; Wen et al. 2024), or by investigating nuclear body-associated RNAs through specific protein labeling techniques (Y. Chen et al. 2018; Fazal et al. 2019).

While these methodologies have contributed novel insights, each approach has its limitations. Methods focusing on RNA-centric interactomes can identify co-binding molecules and binding region sequences but lack information on the genomic locations of these interactions. Proximity-ligation-based methods enable high-throughput genome-wide mapping of RNA footprints on chromatin but are susceptible to capturing numerous non-specific interactions, including those involving diffusive nascent transcripts.

To address these limitations and focus on potentially functional RNA-chromatin interactions, we have developed an approach that combines ribonuclease (RNase) treatment with the in situ mapping of RNA-genome interactome (iMARGI). iMARGI systematically reveals chromatin-associated RNAs (caRNAs) and the genomic regions associated with each caRNA (Wu et al. 2019; Yan et al. 2019). In this study, we incorporated RNase A treatment before the standard iMARGI procedure to remove promiscuous, unprotected RNA transcripts and selectively enrich for RNA-inaccessible, potentially functional RNA-chromatin interactions.

By reanalyzing a previously published iMARGI dataset with this modified protocol (Calandrelli et al. 2023), we provide a comprehensive map of the RNA-inaccessible RNA-chromatin interactome in human embryonic stem cells. Our results reveal that RNA-inaccessible chromatin-associated RNAs (RI-caRNAs) exhibit high sequence conservation and preferential localization to functional genomic regions, including promoters, transcription factor binding sites, and histone modifications. We identify specific RNA classes associated with these regulatory genomic regions, offering new insights into the functional roles of nuclear RNAs in gene regulation and chromatin organization.

## Results

### Overview of RNase-treated iMARGI libraries

In this study, we reanalyzed previously published RNase-treated iMARGI datasets to investigate the non-diffusive RNA-chromatin interaction profile genome-wide (Figure 1a). The original approach involved perturbing the cell nucleus with RNase A (an endoribonuclease that targets single-stranded RNA) before crosslinking, thereby eliminating unprotected RNA from the nucleoplasm and chromatin, and enriching for non-diffusive interactions. Compared to the standard iMARGI protocol, the RNase-treated method included a 10-minute RNase A treatment before crosslinking, followed by the standard iMARGI protocol (Methods). For this analysis, three biological replicates were generated for the RNase-treated samples, and five technical replicates were prepared using the unmodified iMARGI protocol (control libraries) in the human embryonic cell line (H1). In total, 1,589,053,749 and 2,485,334,610 mappable non-duplicated read pairs were generated for the RNase-treated and control libraries, respectively (Supplementary Table 1). We selected read pairs with both RNA and DNA ends uniquely mapped to the human genome and with a mapping score (MAPQ) greater than 30 for downstream analysis to ensure high confidence and quality; these pairs are referred to as valid pairs hereafter. These stringent criteria resulted in 192,077,439 usable read pairs in the three RNase-treated libraries combined and 341,486,278 in the five control libraries combined.

**Figure 1.**
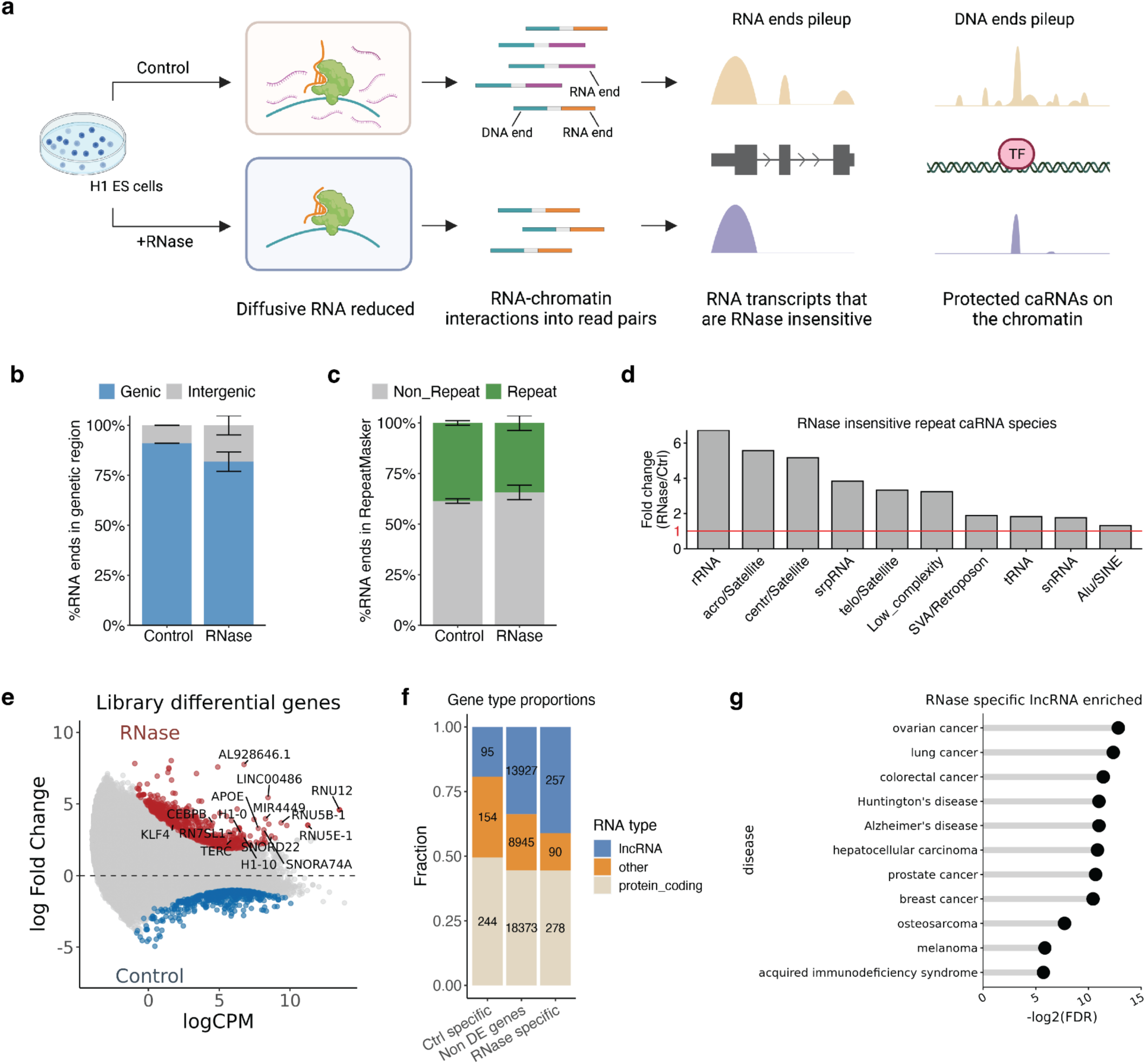
Overview of RNase-treated iMARGI data in H1 ES cells. a) Schematic comparison of standard iMARGI (control) and RNase-treated iMARGI protocols, illustrating differences in RNA-chromatin interactions, RNA transcript profiles, and protected chromatin-associated RNA (caRNA) interacting genomic regions. b) Percentage of RNA reads mapping to genic (blue) and intergenic (gray) regions in control and RNase-treated iMARGI samples. Error bars represent standard error across replicates. c) Percentage of RNA reads mapping to repeat (green) and non-repeat (gray) regions in control and RNase-treated iMARGI samples. d) Fold change of RNase-insensitive repeat caRNA species compared to control. Red line indicates no change (fold change = 1). e) Differential gene expression analysis between control and RNase-treated samples. Red and blue dots represent significantly up- and down-regulated genes, respectively. f) Proportions of lncRNA, protein-coding, and other RNA types among control-specific, non-differentially expressed, and RNase-specific genes. g) Diseases associated with RNase-specific lncRNA, ranked by statistical significance (-log2(FDR)).

Library reproducibility was accessed using RNA ends or DNA ends from iMARGI read pairs individually. We first measured the RNA ends reproducibility by gene read counts using RNA ends only. The lowest correlation between any two pairs of RT-libraries is 0.908 (spearman correlation, p value<2.2e-16) and is 0.928 between any pairs of control libraries (Supplementary Figure 1a, b). For DNA ends, we calculated the genome-wide chromatin attachment levels using sliding windows of 100 kb and also observed high correlations between replicates, although slightly lower than those for RNA ends (Supplementary Figure 1c, d). These data suggest that the RNase-treated iMARGI libraries are highly reproducible.

### Long distance RNA-chromatin interactions are resistant to RNase treatment

An important aspect of RNA-chromatin interactions is the genomic distance between the RNA transcription site and its interacting chromatin regions. Different interaction distances are likely to be governed by distinct mechanisms. For example, extremely short-distance interactions might be attributed to local diffusion of nascent transcripts, while long-distance interactions are more likely to be mediated by RNA-binding proteins (Li and Fu 2019). We first investigated whether the distribution of genomic distances between RNA and DNA ends was altered after RNase A treatment. To this end, we calculated the genomic distance between the RNA and DNA ends for every valid read pair. We found that short-distance interactions (RNA-DNA end distance < 1 kb) were selectively preserved, increasing from 23.00% of total pairs in the control library to 84.53% in the RNase library (Supplementary Figure 2a, proportion test: p-value < 2.22e-16). However, for read pairs with RNA-DNA end distances longer than 1 kb, a larger proportion of distal pairs (>2Mbp or interchromosomal) were significantly enriched (Supplementary Figure 2b, c, proportion test: p-value < 2.22e-16). The selective preservation of long-distance RNA-chromatin interactions was independent of individual chromosomes. These results suggest that chromatin-associated RNAs (caRNAs) involved in short-to middle-range interactions are more susceptible to RNase treatment. The enrichment of short-distance RNA-genome interactions appears RNase resistant, potentially due to protection by RNA polymerases. In contrast, RI-caRNAs participating in distant interactions (RNA-DNA end distance >1 kb) may be protected by other chromatin-binding proteins and are the focus of our subsequent results.

### RNase-inaccessible caRNA transcripts characteristics

We profiled RI-caRNAs features by examining RNA ends from iMARGI read pairs. 83.58% of RNA ends mapped to genes (GENECODE V36 (Frankish et al. 2019), strand specific) in the RNase-treated library, compared to 90.94% in the control (Figure 1b), suggesting genic caRNAs are more accessible to RNase. 33.19% and 37.42% of caRNAs were derived from repeat elements in RNase and control respectively (Figure 1c). While overall repeat-derived caRNA proportions were similar, some classes like rRNA, satellite repeats, and low complexity RNAs were significantly enriched in the RNase library (FCs: 6.73, 5.17, 3.32, 3.24) (Figure 1d), suggesting a subset of intergenic and repeat caRNAs are RNase-inaccessible (Thakur and Henikoff 2020).

Among all genic transcripts, we asked whether there are any genes whose transcripts are inaccessible after RNase A treatment. We called 624 RNase specific genes and 493 control library specific genes using differential analysis (Figure 1e, Methods). We first noticed that lncRNAs (n=257) are over-represented among these 624 RNase specific genes, accounting for 41.93% in RNase library compared to 19.27% in Control library specific genes (pvalue < 2.22e-16, Fisher exact test. Figure 1f). This result suggests that lncRNA transcripts are preferentially protected from RNase A digestion. To link the identified RNase-specific genes with known functions such as disease genesis, we annotated all RNase-specific genes using the DisGeNet database, which includes 20,158 genes associated with 21,775 diseases (Piñero et al. 2020). Similarly, we annotated all lncRNAs with the LncRNADisease database, which includes 19,485 genes associated with 498 diseases (G. Chen et al. 2013) (Figure 1g). The intersections between both libraries are significant (Supplementary Figure 2d, chi-square test, pvalue<2.2e-16). Disease ontology enrichment test also suggests all RNase specific lncRNAs enriched with disease terms including several cancers, Alzheimer’s disease, Huntington’s disease and acquired immunodeficiency syndrome and others (hypergeometric test, FDR<0.05, Supplementary Figure 2e, Methods). In summary, these results suggest that RNase specific genes are enriched with lncRNAs and significantly associated with diseases.

### RNase-inaccessible caRNAs have high level of evolutionary conserveness

Previous studies using RNase-mediated protein footprint sequencing identified protein-protected RNA–protein interaction sites with conserved motif sequences (Ji et al. 2016). We hypothesized that the RNA ends inaccessible to RNase are protected by proteins, and thus are evolutionarily conserved. To test this, we calculated conservation scores of all 192 million RNA ends in the RNase-treated iMARGI library and 341 million RNA ends in the control library. We used phaseCons100 (multiple alignment of 100 vertebrate genomes) (Kent et al. 2002) as a measure of sequence conservation level. We categorized all pairs into three groups based on their RNA-DNA end distance: 1) short-distance pairs (<1kb), 2) cis pairs (>1kb - 2Mb), 3) trans pairs (>2Mb and interchromosomal). We calculated the conservation score at single base resolution, ranging from the center of each read to 500 bp flanking regions. The conservation level was consistently higher for RNA ends in the RNase library compared to control in every group (Figure 2a, b: dashed lines vs. solid lines, t-test p-value < 2.22e-16). Trans pairs exhibited higher conservation than cis pairs, while short-distance pairs had the lowest scores. These results suggest the RNase inaccessible caRNA is more conserved than the other caRNAs.

**Figure 2.**
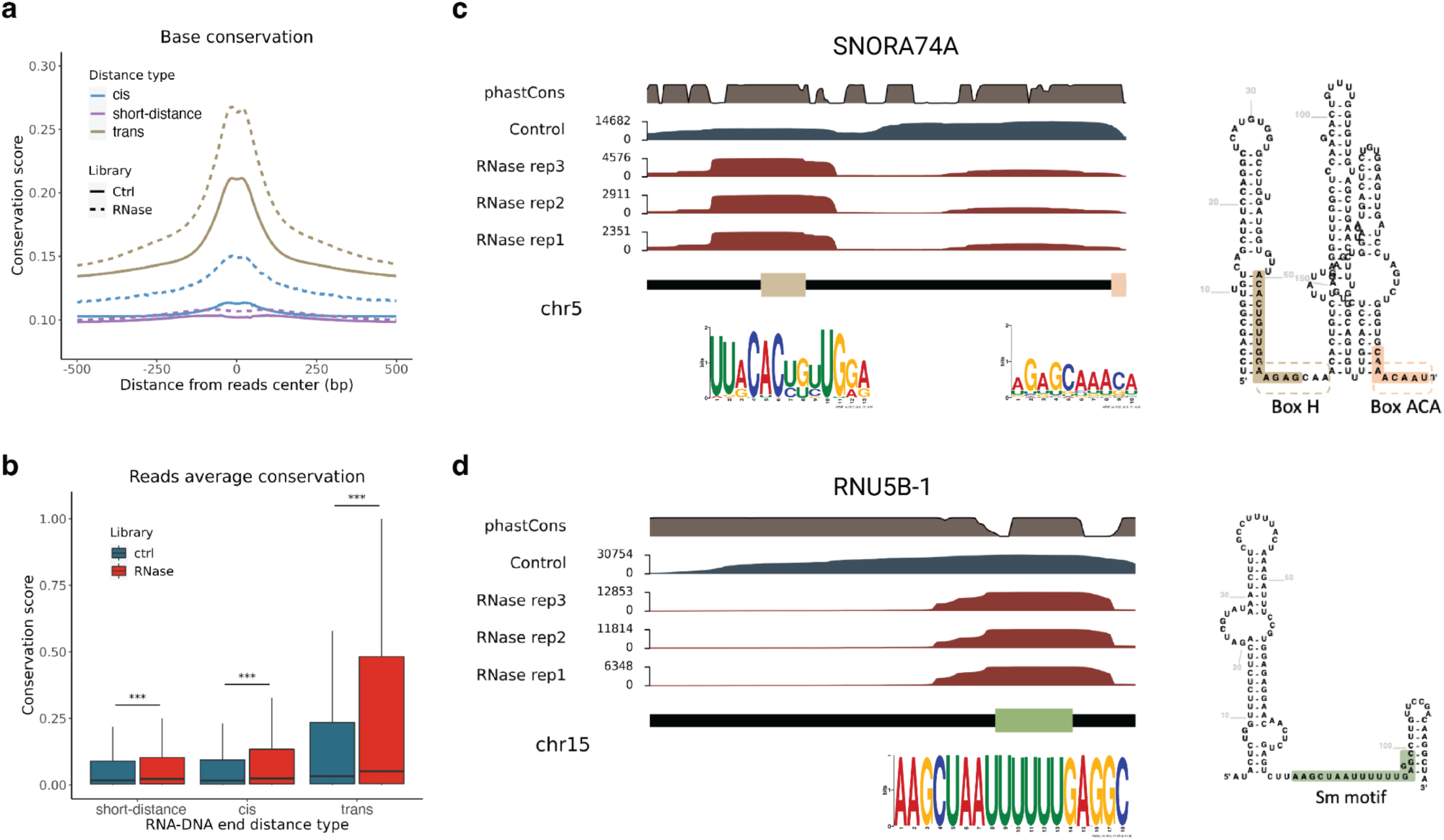
Sequence conservation of RNase inaccessible RNA ends. a) Sequence conservation (PhaseCons 100 score) of each read’s 1000 bp flanking region. All pairs in the control and RNase library were divided into three groups based on their RNA and DNA end genomic distance. short-distance pairs (<1k bp), cis pairs (1k bp ∼ 2Mb), trans pairs (>2Mb and interchromosomal). b) Average read conservation score in each library and each group. Differences of conservation score in each group were tested using t-test. c) RNA ends coverage of SNORA74A gene in three replicates of RNase libraries and one combined control library. Sequence conservation score phastCons100 were shown on the top track. Higher value indicates higher level of conserveness. Secondary structure of SNORA74A was computed by R2DT. RNA end reads enriched regions were highlighted on the secondary structure. Enriched binding sequence motif was called by MEME. d) Same information as c) for gene RNU5B-1. Green box corresponds to the Sm motif.

To gain a clearer picture of what the remaining RNA ends after RNase digestion look like, we analyzed the reads coverage and sequence features of two non-coding genes from the top enriched RNase library specific RNAs: snRNA RNU5B-1 and snoRNA SNORA74A. Reads covering RNU5B-1 are piled up at the Sm domain at the tail of the gene (Figure 2d). Sm domain is a conserved region featured by the consensus sequence Purine-AU4-6G-Purine, and is recognized and bound by spliceosomal Sm proteins (Urlaub et al. 2001). The top one sequence motif identified from all RNA ends transcribed from RNU5B1 is consistent with Sm motif (Figure 2d). Reads transcribed from SNORA74 are overlapped with snoRNA featured H/ACA box motif (Figure 2c) (Urlaub et al. 2001). These examples illustrate that RNaseA treatment was less effective at removing transcripts within the functional regions of a gene whose parts are likely to be interacted with proteins.

### Majority of RNase-inaccessible caRNAs targeting regions are functional genomic regions

To understand the potential functional roles of RNase-inaccessible caRNAs, we focused our analyses on RNA-DNA interaction pairs with RNA and DNA end genomic distances longer than 1 kbp. We first investigated their localization patterns on the genome by calling peaks using DNA end coverage across the genome. In total, 99,784 peaks were identified in the RNase library, with a fragments in peaks ratio (FRiP) of 13.15%, indicating a high signal-to-noise ratio (ENCODE uses FRiP > 1% to filter high-quality transcription factor ChIP-seq experiments (Landt et al. 2012)). After applying the peak max depth threshold of 15 reads, 25,333 peaks, referred to as RNA Attachment Hot zones (RAHs), were identified for further analysis.

Around 49.4% of RAHs were in intronic regions, followed by 23% in intergenic and 16.8% in promoter regions. To assess significance, we compared these proportions to 2,062,460 transcription factor binding sites (TFBS) from 71 TFs in H1 cells (Zheng et al. 2019) and randomized genomic regions with the same length distribution as RAHs. The 16.8% RAH overlap with promoters was significantly higher than 6.26% for randomized regions (permutation test, p-value < 0.001, n=1000), but lower than the 28.69% TFBS promoter overlap (Figure 3d). While the 49.37% intronic RAH proportion was large, it was comparable to the 47.53% genomic background. The 22.97% intergenic RAH proportion was approximately half of the 40.19% background. These data suggest promoter regions are hotspots for caRNA.

**Figure 3.**
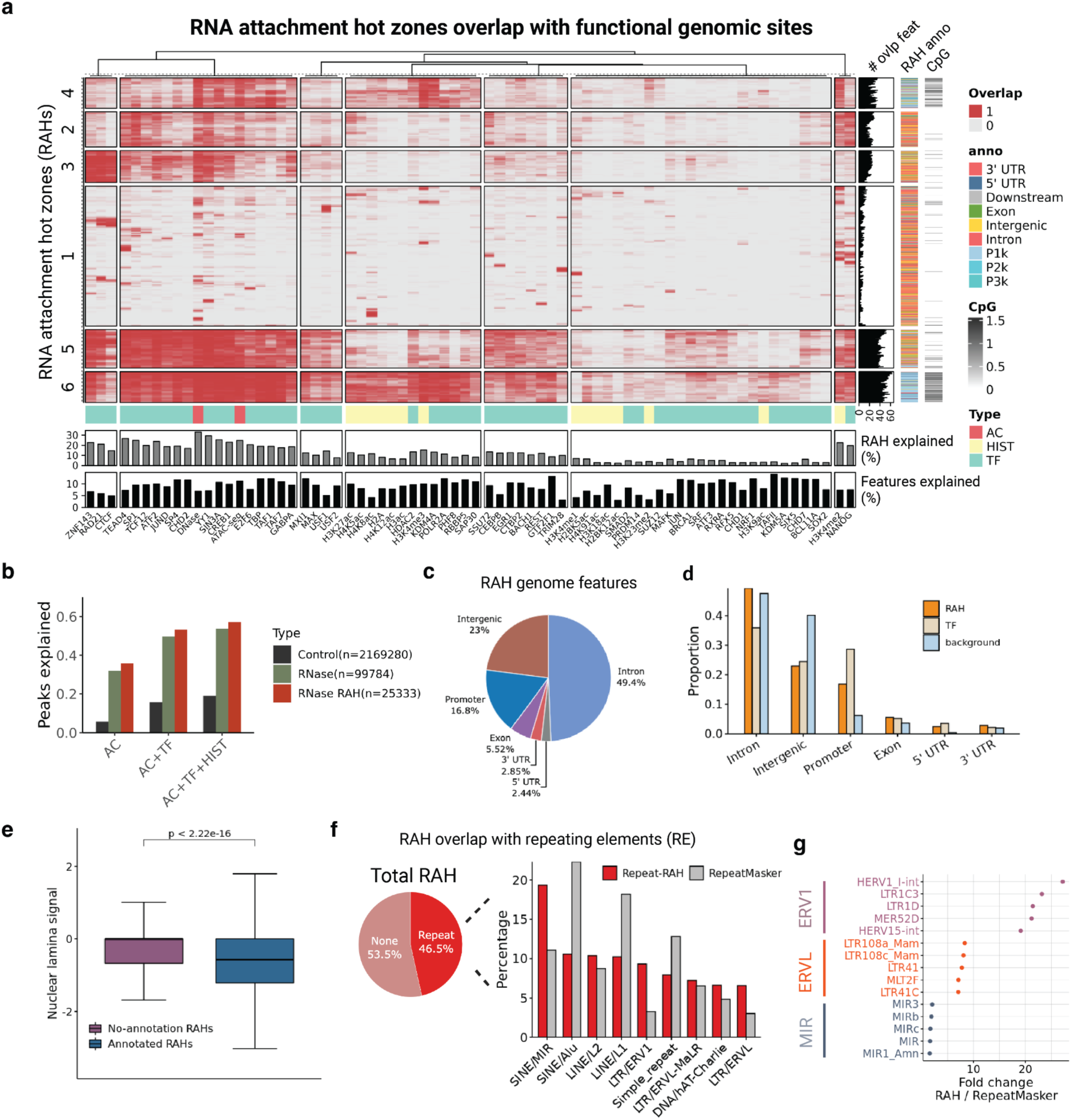
RNAs interacting with functional genomic regions genome wide. a) Heatmap showing overlap between RAHs (n=25,333) and various genomic features (transcription factor binding sites, histone modifications, chromatin accessibility). Rows represent individual RAHs, columns represent genomic features. Color intensity indicates degree of overlap. Tracks below show feature type, proportion of RAHs overlapping each feature, and proportion of feature peaks overlapping RAHs. Right-side tracks display a number of features overlapping each RAH, genomic location, and CpG content. b) Proportion of DNA end peaks explained by chromatin accessibility (AC), transcription factor binding (TF), and histone modifications (HIST) in control, RNase, and RNase RAH conditions. c) Distribution of RAHs across different genomic features. d) Comparison of genomic feature composition between RAHs, transcription factor binding sites, and background genomic regions. e) Nuclear Dam-LaminB1 signals in RAHs with (blue) and without known functional annotations (purple). f) Proportion of RAHs overlapping with repeat elements (RE) and their distribution across RE categories. g) Top 5 repeat element species showing highest fold-change enrichment in RAHs compared to genome-wide RepeatMasker proportions.

We then annotated RAHs with chromatin features: 1) Accessibility (ATAC-seq and DNase I hypersensitive sites), 2) Histone modifications, 3) TFBS, 4) Nuclear periphery (Figure 3a). 35.88% of RAHs overlapped open chromatin regions marked by DNase I or ATAC-seq in H1 cells, six-fold higher than 5.8% of 2,169,280 control library peaks (Figure 3b). This RNA targeting preference mirrors transcription factors preferring open chromatin (ENCODE). Additionally, 53.85% of RAHs overlapped with one or more of the TFBS from the 71 TFs surveyed. Histone modifications further increased the overlap ratio with RAH to 63.71% (Figure 3b). In contrast, only 29% of control peaks were explained by open chromatin, TFBS, and histone modifications (Figure 3b). These data suggest RNase-inaccessible caRNA co-localize to the functional genomic regions.

Some individual transcription factors (TFs) extensively overlap with RAHs. For example, 29.50% of all RAHs overlapped with YY1 binding sites, 26.77% with TEAD4 binding sites, and 14.64% with CTCF. Reciprocally, RAHs also accounted for a sizable proportion of individual TF ChIP-seq peaks. Ranking TFs based on the proportion of their TFBS overlapping with RAHs, we found EZH2 (18.60%), SUZ12 (17.53%), POU5F1 (14.42%), TAFII (14.05%), SP2 (13.91%), and others. All ratios were significantly larger than randomized genomic regions. Interestingly, top-ranked TFs with TFBS explained by RAHs have emerging evidence suggesting dual roles as transcription factors and RNA-binding proteins (see discussion). RAHs simultaneously overlapping multiple TFBS were highly enriched in promoter regions and CpG-rich regions (last two columns, Figure 3a). Notably, 7,551 RAHs (28.90%) overlapped with histone variant H2A.Z peaks (Figure 3a, column cluster 9), the largest factor explaining RAHs after DNase open chromatin regions (33.70%). These results might indicate potential interplay between RNA molecules and the H2A.Z variant. Hierarchical clustering of the RAH-feature overlap matrix highlighted functionally similar markers with similar RAH overlap profiles, e.g., cohesion/insulator markers CTCF, RAD21, and ZNF143 clustered in column cluster 6 (Figure 3a). Polycomb complex histone marker H3K27me3, EZH2, and SUZ12 clustered in column cluster 1 (Figure 3a).

For 8,400 RAHs not overlapping known TFBS or histone modifications, we asked whether they were enriched in inaccessible regions like heterochromatin and the nuclear lamina. We utilized Lamin B1 DamID signals, reflecting a genomic region’s proximity to the nuclear lamina (Guelen et al. 2008), to annotate each RAH. RAHs without TF binding, chromatin accessibility, or histone modification annotations had significantly higher DamID-Lamin B1 signals, suggesting closer physical proximity to the nuclear lamina and heterochromatin (t-test, p-value < 2.2e-16, Figure 3e).

Finally, we asked whether RAHs overlapped with any repetitive elements (REs). Only 46.5% of RAHs overlapped with UCSC RepeatMasker-annotated REs, less than the 68.31% background from randomized regions with the same length distribution as RAHs (Figure 3f), suggesting RAHs are less enriched in genomic RE regions. Among RE-overlapping RAHs, SINE/MIR, LTR/ERV1, and LTR/ERVL were three RE classes significantly associated with RNase-insensitive caRNAs, with 1.74-, 2.85-, and 2.71-fold enrichment compared to the proportion of genome containing each repeat class annotated in RepeatMasker annotations (Figure 3g). Interestingly, previous studies suggested SINE/MIR (mammalian-wide interspersed repeats) are a conserved transposable element family with significant regulatory capabilities and sequence similarities to tRNA-related genomic insulators (Pol III binding), independent of the CCCTC-binding factor (CTCF) insulation mechanism (Wang et al. 2015). Similarly, the LTR/ERV1 and LTR/ERVL families include HERV-int repeats proposed to demarcate topologically associating domains in human pluripotent stem cells (Zhang et al. 2019). Together, these results suggest RI-caRNAs preferentially interact with genomic repeats functioning as organizational insulators.

In summary, these data show RNase-inaccessible caRNA tends to localize at functional genomic regions. Major caRNA localization hotspots include gene promoter regions, transcription factor binding sites, H2A.Z chromatic regions, as well as H3K27me3 modified and Polycomb associated chromatic regions.

### RNase removes promiscuous RNAs and magnifies the stringent interactions between RNA-TFs and RNA-histone modifications

Our previous results indicated that chromatin-associated RNA (caRNA) genome interaction hot zones (RAHs) frequently colocalize with transcription factor binding sites (TFBS) or histone modifications. To further investigate this relationship, we conducted a comprehensive analysis of caRNA co-localization with TFBS and histone modifications using all ChIP-seq data from the Cistrome and ENCODE databases (ENCODE Project Consortium 2012; Liu et al. 2011) for the H1 cell line.

We assessed caRNA enrichment by comparing the abundance of caRNAs interacting at reference peaks (TFBS or histone modification sites) versus surrounding regions. For each reference marker, we generated heatmaps depicting caRNA coverage in both control and RNase-treated libraries. These heatmaps represent all ChIP-seq peaks from the databases (rows), with columns showing 10kb regions surrounding the peak center (1000 bins). Based on the caRNA-peak abundance patterns in control and RNase libraries, we classified TFs and histone modifications into two categories: Class 1) those with increased caRNA peak center versus background ratio in the RNase library compared to the control, and Class 2) those with decreased ratios.

For Class 1 markers, we observed concentrated caRNA signals at the TFBS ChIP-seq peak centers in both control and RNase libraries (Figure 4a, Supplementary Figure 3a, c). Importantly, the RNase-treated library retained only the TFBS-associated RNAs, while depleting the surrounding signals (Figure 4c). To quantify this enrichment, we developed a concentration score, calculated as the ratio of caRNA coverage between the reference peak center (±500bp) and the surrounding region (±1500bp). Higher scores indicate stronger enrichment at the central peak. The uniformly distributed caRNA coverage would yield a concentration score at 0.3.

**Figure 4.**
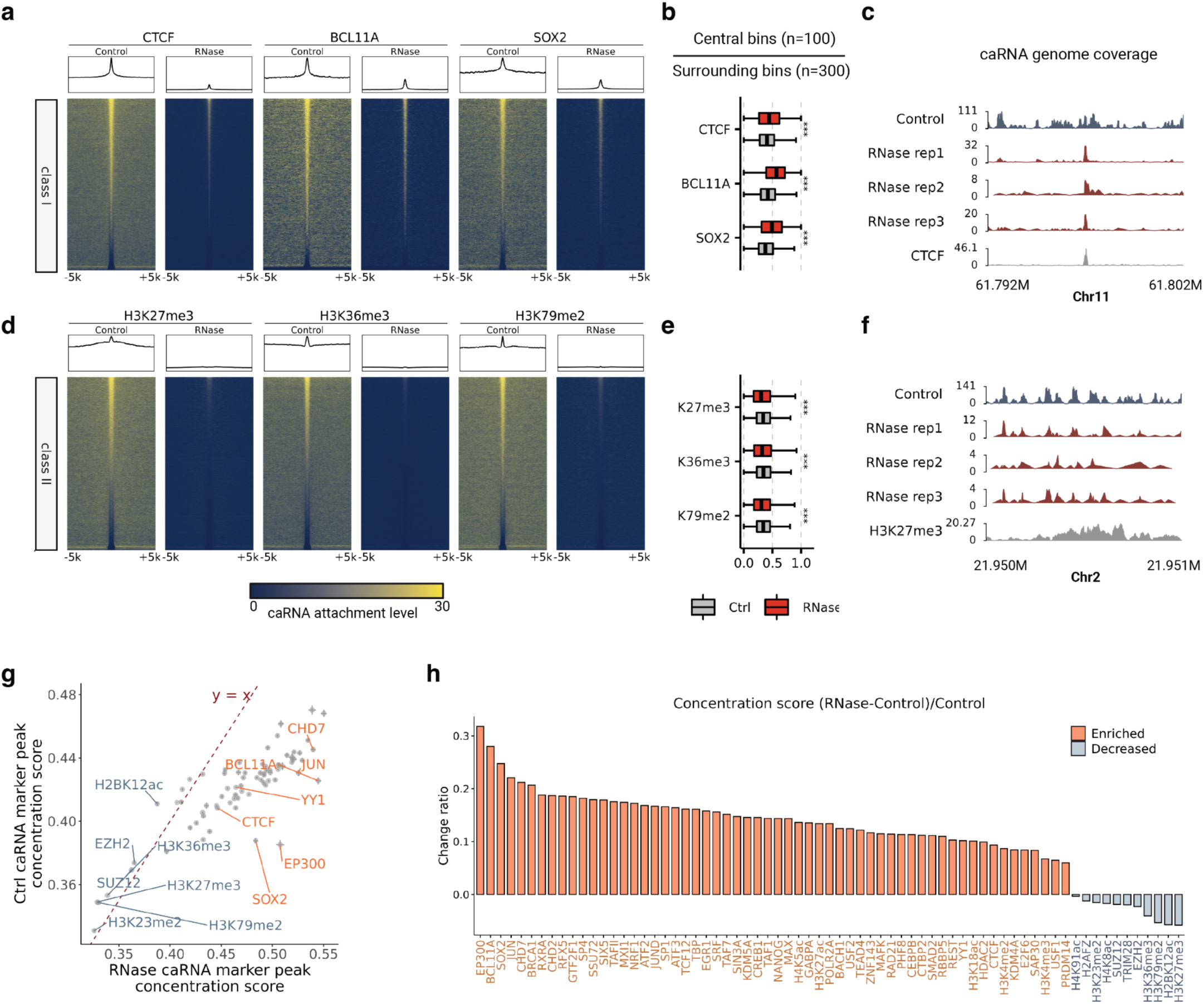
RNase treatment magnifies stringent interaction of caRNA-TF interactions. a) Heatmaps and summary plots for CTCF, BCL11A, and SOX2 binding sites. Each row represents a ChIP-seq peak (peak center ±5kb) from H1 cells. Color intensity indicates caRNA interaction level. Class 1 transcription factors show enhanced caRNA-TF peak interactions after RNase treatment. b) Boxplots comparing caRNA concentration scores in control vs RNase-treated samples for the transcription factors in (a). Concentration score is the ratio of caRNA attachment in the central 1kb to the surrounding 3kb of each peak. c) Genome browser view of caRNA attachment levels at a CTCF binding site on Chr11. Tracks show control and three RNase-treated replicates. d-f) Similar to (a-c), but for histone modifications H3K27me3, H3K36me3, and H3K79me2. Class 3 histone modifications show diminished caRNA signals after RNase treatment. g) Scatter plot comparing caRNA concentration scores between control and RNase-treated samples for 76 chromatin markers. Dashed line represents y=x. h) Bar plot showing changes in concentration scores ((RNase-Control)/Control) for 69 invariant chromatin markers. Orange and blue indicate enriched and decreased scores, respectively.

The median concentration score for CTCF, a representative Class 1 marker, increased from 0.405 in the control library to 0.445 in the RNase library, indicating specific targeting of CTCF peaks by caRNAs. Paired t-tests confirmed significantly higher concentration in the RNase library for each CTCF peak (Figure 4b). We observed similar trends for other Class 1 TFs, including BCL11A (0.429:0.570), EP300 (0.374:0.523), SOX2 (0.379:0.492), RAD21 (0.408:0.462), CHD7 (0.441:0.559), and YY1 (0.419:0.468) (Figure 4b, Supplementary Figure 4a, b). Paired t-tests for all above TFs showed significant enrichment (p-value < 2.2e-16). These results demonstrate that RNase treatment does not uniformly decrease RNA abundance but selectively preserves RNA signals at TF binding regions. The RNA signals at the TF binding regions are selectively preserved. These data are in line with recent reports on RNA’s roles in facilitating protein-chromatin binding (Hansen et al. 2019; Saldaña-Meyer et al. 2019; Sigova et al. 2015; Xiao et al. 2019).

In contrast, Class 2 markers showed significant depletion of caRNAs at their associated regions after RNase treatment. This category includes suppressive marks like H3K27me3 (control:RNase, 0.337:0.305, p < 2.2e-16), gene body histone mark H3K36me3 (0.342:0.320, p < 2.2e-16), and active gene histone mark H3K79me2 (0.340:0.310, p < 2.2e-16) (Figure 4d-f). These histone modifications typically form broad peaks and require continuous deposition through protein complexes to maintain their states. The attenuated peaks in RNase-treated samples suggest that caRNAs co-localizing with these histone marks are not protected by stable protein associations. Decreased caRNA levels have also been observed for class II markers such as histone modifications H3K23me2 (0.320:0.298), H2BK12ac (0.411:0.373), TRIM28 (0.419:0.411), EZH2 (0.373:0.365), and SUZ12 (0.369:0.362).

To ensure the robustness of our findings, we tested different combinations of background and peak central bin numbers for calculating concentration scores. This comprehensive analysis covered all histone modification ChIP-seq and TFBS data available from Cistrome and ENCODE for the H1 cell line (Figure 4g). In total, 69 markers showed consistent significant changes across various concentration calculation metrics, with 58 markers displaying enriched caRNA levels at their peaks after RNase A digestion and 11 markers showing decreases (Figure 4h).

Intriguingly, literature analysis revealed that 22 out of the 58 iMARGI-identified Class 1 TFs are known to be both RNA and DNA-binding proteins. This group includes EP300, SOX2, JUN, CHD7, BRCA1, CHD2, GTF2F1, ATF2, EGR1, TAF7, MAX, SIN3A, GABPA, POLR2A, USF2, RAD21, CTCF, CEBPB, RBBP5, and YY1, among others. This finding suggests a potential mechanism where caRNAs are protected from RNase digestion through direct protein binding.

In conclusion, our data indicate that caRNA-TF associations are pervasive and apply to numerous TFs. The RT-iMARGI technique magnifies the stringent interactions between TFs and caRNAs that are integral components of the chromatin landscape and robust to RNase perturbation. The attachment profile appears to be highly specific to the intrinsic characteristics of the target, providing new insights into the complex interplay between RNA, proteins, and chromatin in gene regulation.

### Transcription factor specifically associated RNA classes

Building on our previous findings that chromatin-associated RNA molecules (caRNAs) target transcription factor binding sites (TFBS) with RNase-resistant associations, we sought to identify specific RNA species enriched at TFBS for individual transcription factors (TFs).

To identify RNA species enriched in associating with specific TFs, we compared the relative abundance of TFBS-targeting caRNA and all iMARGI detected caRNA. Specifically, we tested the dependence between gene_i targeting TF_j and RNase treatment using a chi-square test. The null hypothesis is that there is no dependence between gene_i targeting TF_j TFBS and RNase treatment, such that the chance of seeing gene_i transcripts targeting all TFBS of TF_j compared to not targeting TF_j TFBS would be no different than without RNase treatment. The alternative hypothesis was a dependence such that RNase treatment selectively preserved some TF-associated caRNAs (Figure 5a). We tested TF-enriched genes for each TF that had increased center to flanking region caRNA attachment level in the RNase library from the previous section, examining 52 TFs. The chi-square p-value distributions were all heavily skewed towards 0, indicating TF-associated genes significantly enriched in the RNase-treated library (Supplementary Figure 4a). We identified thousands of genes enriched for each TF under an adjusted p-value < 0.05 (Benjamini & Yekutieli 2001, no assumption on gene independence) and odds ratio > 1 (Supplementary Figure 4b).

**Figure 5.**
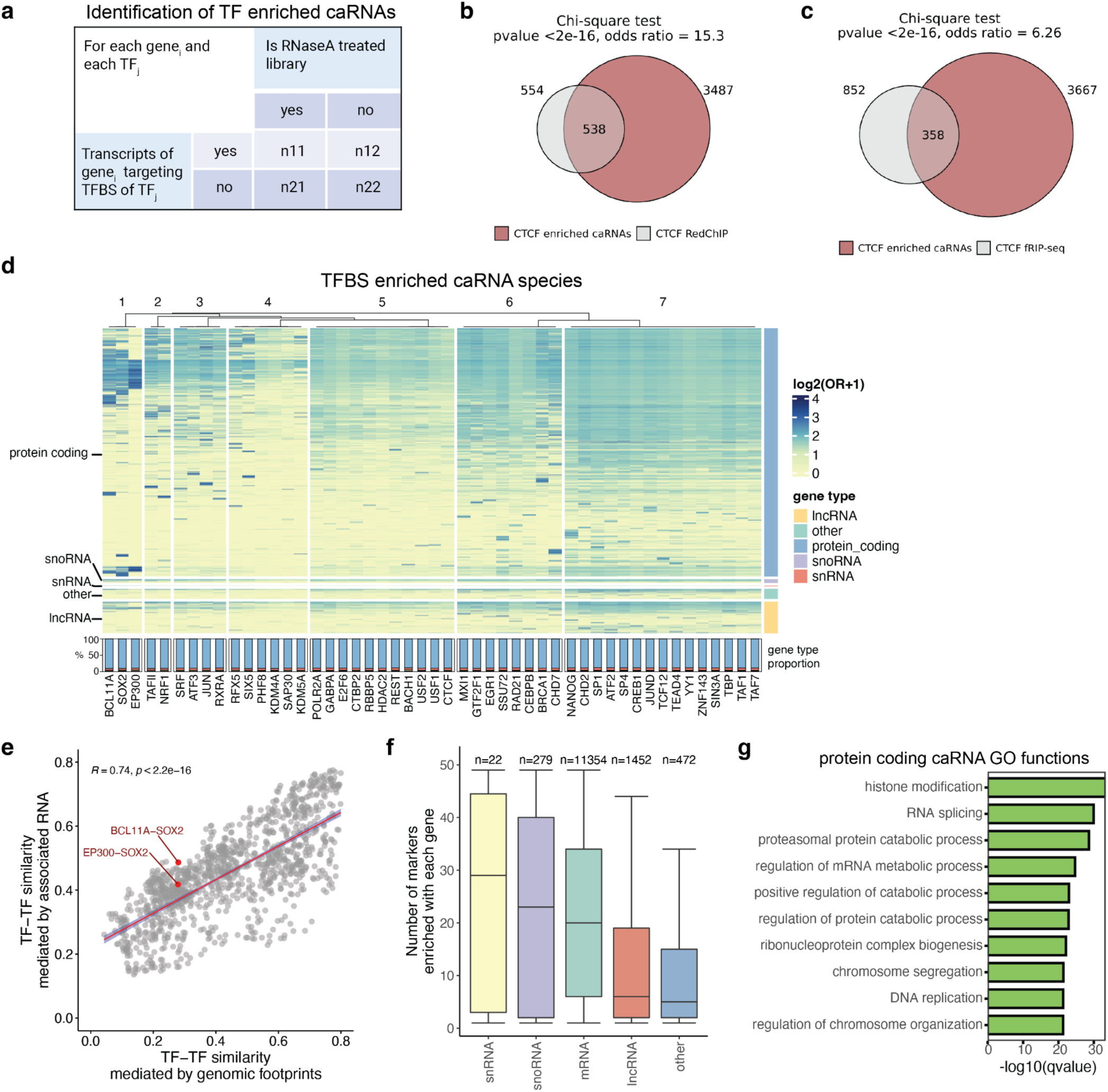
Transcription factor binding sites enriched caRNA species. a) Contingency table of association test to identify TFBS enriched caRNA species. b) Venn plot showing the overlap between CTCF enriched caRNAs (in H1 cells) and RedChIP identified CTCF associated caRNAs (in K562 cells). c) Venn plot showing the overlap between CTCF enriched caRNAs (in H1 cells) and fRIP-seq identified CTCF associated caRNAs (in K562 cells). d) Heatmap showing each TF enriched caRNAs with color mapped to odds ratio from association test. Columns are clustered using hierarchical clustering based on Jaccard distance. Rows are genes split by gene type. Bottom track showing the proportion of each RNA species among all enriched caRNAs for each TF. e) Correlation between TF-TF similarities measured by TF genome footprints (number of TFBS in each non-overlapped 100kbp bin on the genome) or enriched RNA species. f) Boxplot showing the number of TFs each RNA species associated with. g) Top enriched GO terms of all mRNAs that are enriched with any TF examined.

To examine whether iMARGI-identified TF-specific associated RNA species were expected, we compared the candidate list with two related datasets: 1) RedChIP data, measuring protein-involved chromatin-associated RNAs (Gavrilov et al. 2022), and 2) fRIP-seq data, measuring protein-bound RNA by formaldehyde RNA immunoprecipitation (G Hendrickson et al. 2016). The first dataset offers an orthogonal strategy for detecting caRNAs by pulling down target proteins and detecting associated DNA and RNA. Although the second dataset does not directly measure caRNAs, it aligns with our hypothesis that RNase treatment-insensitive caRNAs may be protected by protein binding. We found that iMARGI-identified CTCF TFBS-enriched genes significantly overlapped with both RedChIP-detected CTCF-associated caRNAs (RedChIP data generated in K562 cells; odds ratio = 15.3, p-value < 2.2e-16) and fRIP-seq-detected CTCF-bound RNAs (fRIP-seq data generated in K562 cells; odds ratio = 6.26, p-value < 2.2e-16) (Figure 5b, c). The overlap with RedChIP was even more significant than fRIP-seq, consistent with the idea that both iMARGI and RedChIP detect chromatin-associated RNA components.

Similarly, TFs CHD7 and RBBP5 were surveyed in both iMARGI (in H1 cells) and fRIP-seq (in K562 cells). fRIP-seq nominated 994 CHD7 protein-bound RNAs, 361 of which overlapped with iMARGI-identified CHD7 binding site-enriched caRNAs (odds ratio = 6.46, p-value < 2.2e-16) (Supplementary Figure 4c, d). For RBBP5 (Retinoblastoma-binding protein 5), fRIP-seq identified 984 specific genes, and 107 overlapped with RBBP5 binding site-enriched caRNAs (odds ratio = 2.12, p-value = 5.4e-15) (Supplementary Figure 4e, f). Together, these data suggest RT-iMARGI-identified CTCF TFBS-enriched caRNAs can be rationalized using orthogonal technologies.

Since iMARGI-derived caRNA-TFBS association specificity does not restrict to any existing antibody of the target protein of interest, we could comprehensively survey all TF-specifically associated caRNAs. We then asked whether any TFs share the same set of associated caRNA genes. We plotted all the caRNA that are enriched in the TFBS of any analyzed TF in a heatmap colored by the degree of association (odds ratio) (Figure 5d). TFs were clustered based on their specifically associated RNA profiles using hierarchical clustering based on Jaccard distance. We found that the RNA-mediated TF-TF similarities are highly correlated with TFBS similarities on the genome, measured by their co-localization (Figure 5e, rho = 0.74, p-value < 2.2e-16, Methods). Nevertheless, some TF pairs showed elevated correlation in RNA-mediated profiles compared to genomic footprints, such as BCL11A, SOX2, and EP300. Recent studies have revealed that the pluripotency factor SOX2 is an RNA-binding protein (Z. E. Holmes et al. 2020) and has direct protein-protein interactions with the coactivator EP300 (B. R. Kim et al. 2017).

For each TF, we annotated the gene species composition (Figure 5d, bottom and right tracks). Surprisingly, protein-coding genes were the largest proportion of gene species for every examined TF. While many non-coding genes have been observed associated with transcription factors and histone marks, the role of protein-coding genes in epigenetic regulation is less studied. We then asked whether any gene species were more likely to associate with many chromatin-associated proteins (CAPs). We found that snRNAs were the most promiscuous type, with the median number of associated markers for all snRNAs being 29, followed by snoRNAs (median = 23), mRNAs (median = 20), and lncRNAs (median = 6) (Figure 5f). We then asked whether the number of associated markers was attributable to high gene expression levels. The Spearman correlation between gene expression and the number of associated markers was 0.671 (Supplementary Figure 4g, p-value < 2.2e-16), suggesting transcripts with higher abundance tend to enrich at more TFBS. These data are consistent with the large proportion of mRNAs observed in both RedChIP and fRIP-seq data, suggesting CAPs bind promiscuous RNAs likely correlated with RNA abundance within the nucleus.

Taken together, these data suggest that different TFs are enriched with different caRNA species that differentiate them from their genomic footprints. Highly abundant gene transcripts, including protein-coding RNAs, are widely observed to be enriched at TFBS.

## Discussion

This study presents an in-depth analysis using an RNase-mediated RNA-chromatin interaction profile to globally investigate the non-diffusive RNA-chromatin interactome genome-wide. Our results reveal several key insights into the nature and function of chromatin-associated RNAs (caRNAs). Historically, when chromatin-associated RNAs were first discovered, the prevailing belief was that they represented a major outcome of nascent transcription, with newly transcribed RNAs attached close to their transcription start sites (D. S. Holmes et al. 1972). Recent studies also suggested that the localization pattern of caRNAs is often determined by their transcription start sites and expression levels (Limouse et al. 2023). However, in other cases, some RNAs can act in trans and play roles at distant genomic sites (Clemson et al. 2009; Tripathi et al. 2010).

Our study demonstrates that long-distance RNA-chromatin interactions are preferentially preserved after RNase treatment, while short-range interactions are depleted. This suggests that caRNAs involved in long-range targeting are more likely to be functionally relevant and protected, potentially by RNA-binding proteins. The enrichment of extremely proximal RNA-genome interactions likely reflects polymerase protection of nascent transcripts. Our findings indicate that RNase treatment can remove the majority of caRNAs contributed by nascent transcripts, while the remaining RNase-inaccessible fraction usually involves long-range interactions.

Our study also demonstrates that RNase insensitive caRNAs exhibit distinct genomic features compared to the total caRNA population. They are enriched for intergenic transcripts, certain repeat classes (e.g. satellites, rRNA), and disease-associated lncRNAs. Furthermore, these RNase-inaccessible caRNAs display greater evolutionary sequence conservation, particularly at known functional RNA domains like Sm sites and H/ACA boxes. This evolutionary constraint implies functional importance for these caRNA molecules. RNase treated iMARGI pinpoints the localization preference of RNase-inaccessible caRNAs to regulatory genomic regions like promoters, transcription factor binding sites (TFBS), and histone variant (H2A.Z) sites. A subset also maps to heterochromatic lamina-associated domains. Integration with epigenomic data suggests interplay between these caRNAs and chromatin regulatory machinery like transcription factors and Polycomb complexes.

RNase-treated iMARGI magnifies the specific associations between caRNAs and regulatory factors like transcription factors. The caRNA binding signals at many transcription factor binding signals are depleted genome-wide after RNase treatment, except at cognate TFBS where caRNA enrichment persists. This concentration effect implies these TFBS-localized caRNAs are protected, potentially by direct binding to the transcription factors themselves. Supporting this, many transcription factors have been suggested to interact with caRNA, such as EP300 (Bose et al. 2017) , SOX2 (Z. E. Holmes et al. 2020), BRCA1 (Vohhodina et al. 2021) and many others. Notably, a recent landmark study (Oksuz et al. 2023) revealed that the ability to bind RNA is a widespread property of transcription factors, with at least half exhibiting direct RNA-binding capabilities mediated by a previously unrecognized arginine-rich RNA-binding domain (ARM-domain). This suggests that the specific caRNA-transcription factor associations observed likely represent a general phenomenon where transcription factors use RNA-binding domains to interact with chromatin-associated RNAs at their regulatory sites across the genome. The ability of RNase-treated iMARGI to pinpoint these specific RNA-protein interactions provides a powerful means to delineate the functional RNA components involved in transcriptional regulation by diverse transcription factors.

In summary, RNase-treated iMARGI enables high-resolution mapping of the functional RNA-chromatin interactome by removing diffusive RNA signals. The RNase-inaccessible interactions pinpoint regulatory RNAs, their genomic localizations, and mechanistic associations with chromatin factors. These findings nominate novel RNA components of nuclear regulation and provide a rich resource for investigating the multi-faceted roles of caRNAs in genome organization and gene expression control. Future studies could leverage RNase-treated iMARGI to probe RNA-chromatin interactions in diverse cellular contexts and link specific RNA molecules to regulatory phenomena. Additionally, integrating RNA perturbations or RNA-binding protein depletions could directly test the functional requirements for caRNAs in chromatin regulation. Overall, this work establishes a powerful approach for deciphering the RNA-centric nodes of nuclear regulation, opening new avenues for understanding the complex interplay between RNA, chromatin, and gene regulation.

## Methods

### Experimental Methods

#### RNaseA treatment

H1 cells were harvested from cell culture plate and aliquoted cell suspension to 10 million H1 cells per 1.5 mL tube. Wash the cells with 1 mL 1X PBS and centrifuge at 500 X g for 3 min at RT. Then, cells were gently permeabilized by resuspending cell pallets with 0.01% PBST (TritonX-100 in PBS) and treated for 5 min at RT. After permeabilization, cells were treated with 200 µg/mL RNase A (Thermo Fisher Scientific, Cat# EN0531) on rotator for 10 min at RT. The treated cells were fixed with 4% formaldehyde (Thermo Fisher Scientific, Cat# 28906) for immunofluorescence imaging. For Hi-C and iMARGI library generation, the treated cells were fixed with 1 mL 1% formaldehyde on the rotator for 10 min at RT. Then, the reactions were terminated with 250 µL 1M glycine on the rotator for 10 min at RT. The treated sample was centrifuged at 2000Xg for 5 min at 4°C and washed with 1 mL cold 1X PBS.

### Quantification and Statistical Analysis

#### Library specific genes identification

We first summarized the number of RNA ends mapped to each gene in control and RNase A treated libraries respectively. We filtered out genes that have zero counts across all replicates. We then applied edgeR to identify the differentially expressed genes between all replicates under the null hypothesis that the fold change of gene abundance between the two conditions is zero. For RNase library the threshold is adjusted p-value cutoff at 10-6, a log fold change increase at 5. For the control library the adjusted p-value cutoff at 0.05, a log fold change decreases at 5. The different thresholds chosen in the two libraries due to the unbalanced expression profile for genes, namely, many genes only have transcripts in the control library.

#### Gene set disease ontology term enrichment analysis

We derived a disease ontology reference by assigning each disease term with their associated genes annotated from the LncRNADisease database. For each of these DE genes related disease-gene sets with size larger than 10 (including more than 10 genes associated with each disease term), we applied the hypergeometric test to see if any disease ontology term is enriched with RNase specific DE genes. Enriched disease terms were selected using FDR < 0.05.

#### RNA attachment hotzones identification

To systematically identify genomic regions enriched with RNA transcripts in the RNase library, we employed the following approach. First, peaks were called using DNA end coverage across the genome to reduce the influence of weak signals. To avoid peaks formed due to nascent transcription, only read pairs with an RNA-DNA end distance greater than 1kb were considered. Peak calling was performed using MACS2 on the combined replicates of the RNase library. The fragments in peaks ratio (FRiP) was calculated to assess the signal-to-noise ratio. To focus on peaks with high read coverage, a peak max depth threshold of 15 was applied, and peaks meeting this criterion were designated as RNA Attachment Hot zones (RAHs).

#### Transcription factor similarities measured by colocalization similarities

TF-TF similarity based on ChIP-seq data localization was calculated as follows: ChIP-seq peak files for each TF were imported as genomic ranges. The human genome was tiled into 100kb bins and for each TF, the number of overlapping peaks per bin was counted, generating a bin x TF matrix. Pairwise TF-TF similarity was then computed as the correlation between the respective rows of this matrix using the *simil* function from the proxy R package.

## Data and Code Availability

RNase treated and Control iMARGI data can be downloaded from 4DN data portal with accession number: 4DNESNOJ7HY7 (Control iMARGI in H1 cell line), 4DNESOBRUQ12 (RNase treated iMARGI in H1 cell line). All custom code used for data analysis in this study will be made publicly available on GitHub upon acceptance of the manuscript.

## Author contributions

X.W. conceived the original idea, designed and performed all data analyses, developed the methodology. X.W., S.Z. wrote the manuscript. S.Z. managed and supervised the project.

---

**Supplementary Figure 1.**
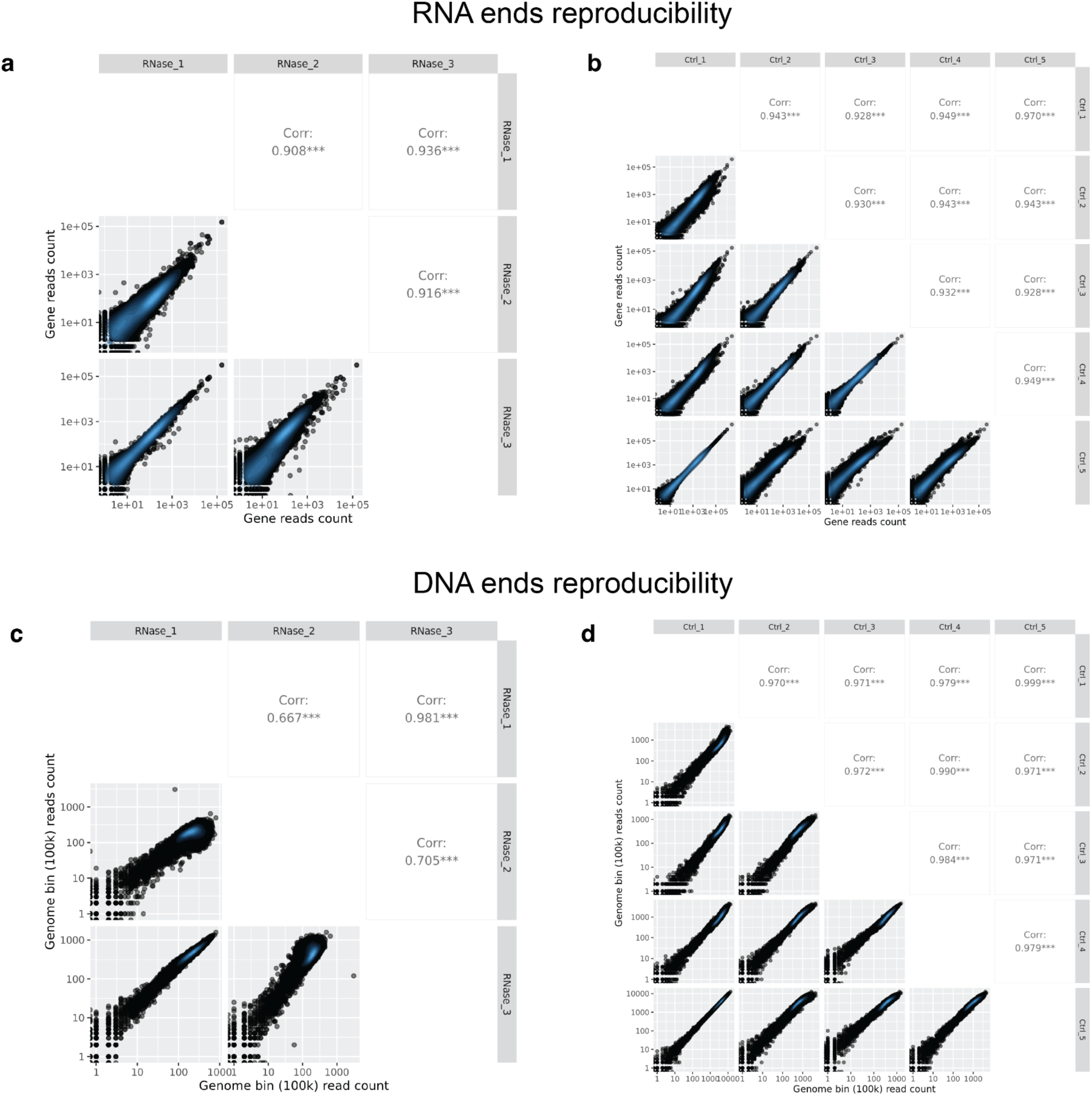
Reproducibility of iMARGI library data for RNA and DNA ends. iMARGI pairs with RNA and DNA end distance larger than 1000bp used. a) Reproducibility of RNA end read counts per gene across three RNase-treated iMARGI replicates. Each point represents a gene, with read counts plotted on a log scale. RNA ends were counted in a strand-specific manner. Spearman correlation coefficients and their associated p-values are shown for each pairwise comparison. b) Similar to (a), but showing reproducibility across five control iMARGI replicates. The matrix of scatter plots illustrates pairwise comparisons between all control samples, with correlation coefficients displayed for each pair. c) Reproducibility of DNA end read counts mapped to 100 kbp genomic bins across three RNase-treated iMARGI replicates. Bins were derived by sliding across the whole genome without overlap. As in (a), Spearman correlation coefficients and p-values are provided for pairwise comparisons. d) Similar to (c), but showing reproducibility of DNA end read counts across five control iMARGI replicates, presented in a matrix of pairwise comparisons with correlation coefficients. p < 0.001, indicated by ***.

**Supplementary Figure 2.**
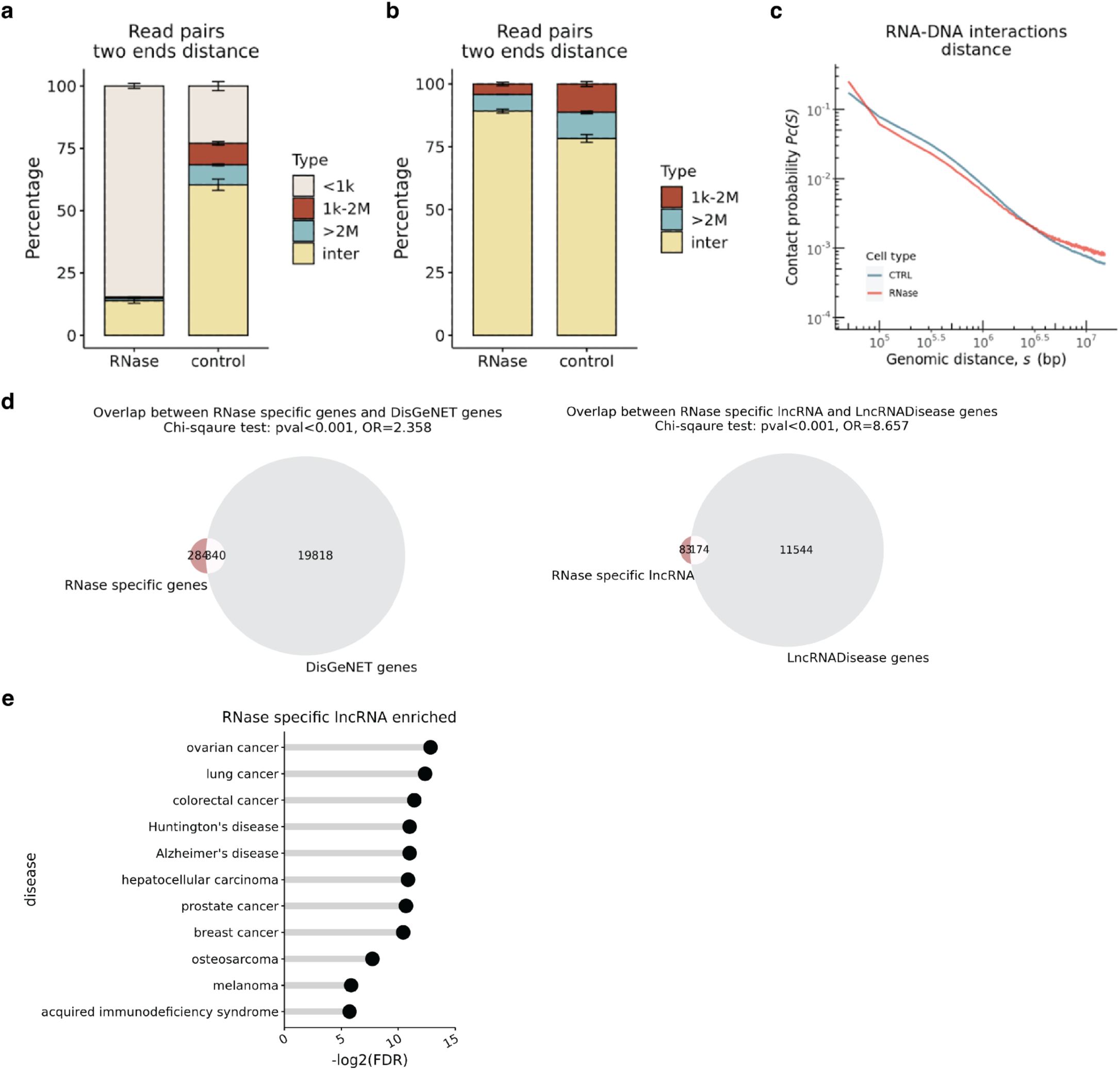
iMARGI RNA, DNA ends distance distribution. a) Barplot showing the proportion of read pairs that with RNA, DNA end distance belongs to <1kbp, 1kbp∼2Mbp, >2Mbp but within one chromosome and finally interchromosomal read pairs. Error bar of the proportion was derived from library replicates. b) Similar to a), but without <1kbp category. c) Genomic distance versus contact frequency for all RNA, DNA read pairs with two ends distance longer than 1kbp. d) Venn plot showing the overlap between RNase treated library specific genes and DisGeNET or LncRNADisease databases. e) RNase specific genes enriched disease terms.

**Supplementary Figure 3.**
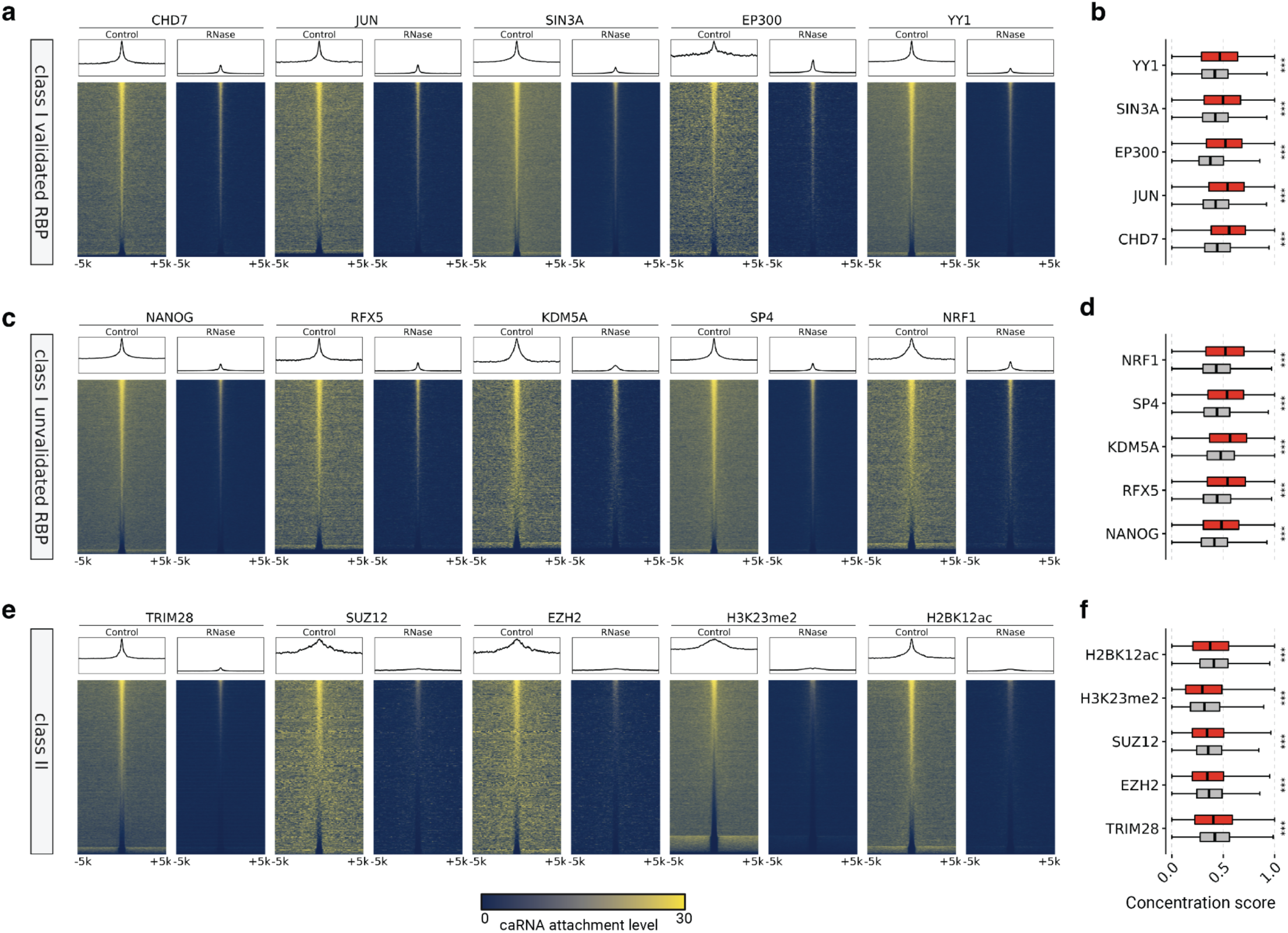
RNase treatment magnifies stringent interaction of caRNA-TF interactions. a) Heatmap and summary plot for five class I TFs (stronger caRNA enrichment at TFBS after RNase digestion) that have been suggested to have RNA binding properties: CHD7, JUN, SIN3A, EP300, YY1. Rows represent all peaks of a TF. Chip-seq data annotated from Cistrome database H1 cell line. Columns include 1000 bins represent 10kb surrounding regions from the center of a chip-seq peak. Color maps to the number of caRNAs interacting with each bin. Summary plot shows the average coverage across all peaks for a transcription factor. b, d, f) Boxplot of concentration score in Control or RNase library. The score is defined as peak central 1000bp region caRNA attachment level divided by peak central 3000bp caRNA attachment level. c) Similar to a), showing five TFs have enriched caRNA binding after RNase digestion but have not been reported has RNA binding property. e) Similar to a), showing five markers including TF TRIM28 or histone modifications whose DNA sites showing depleted caRNA level after RNase digestion.

**Supplementary Figure 4.**
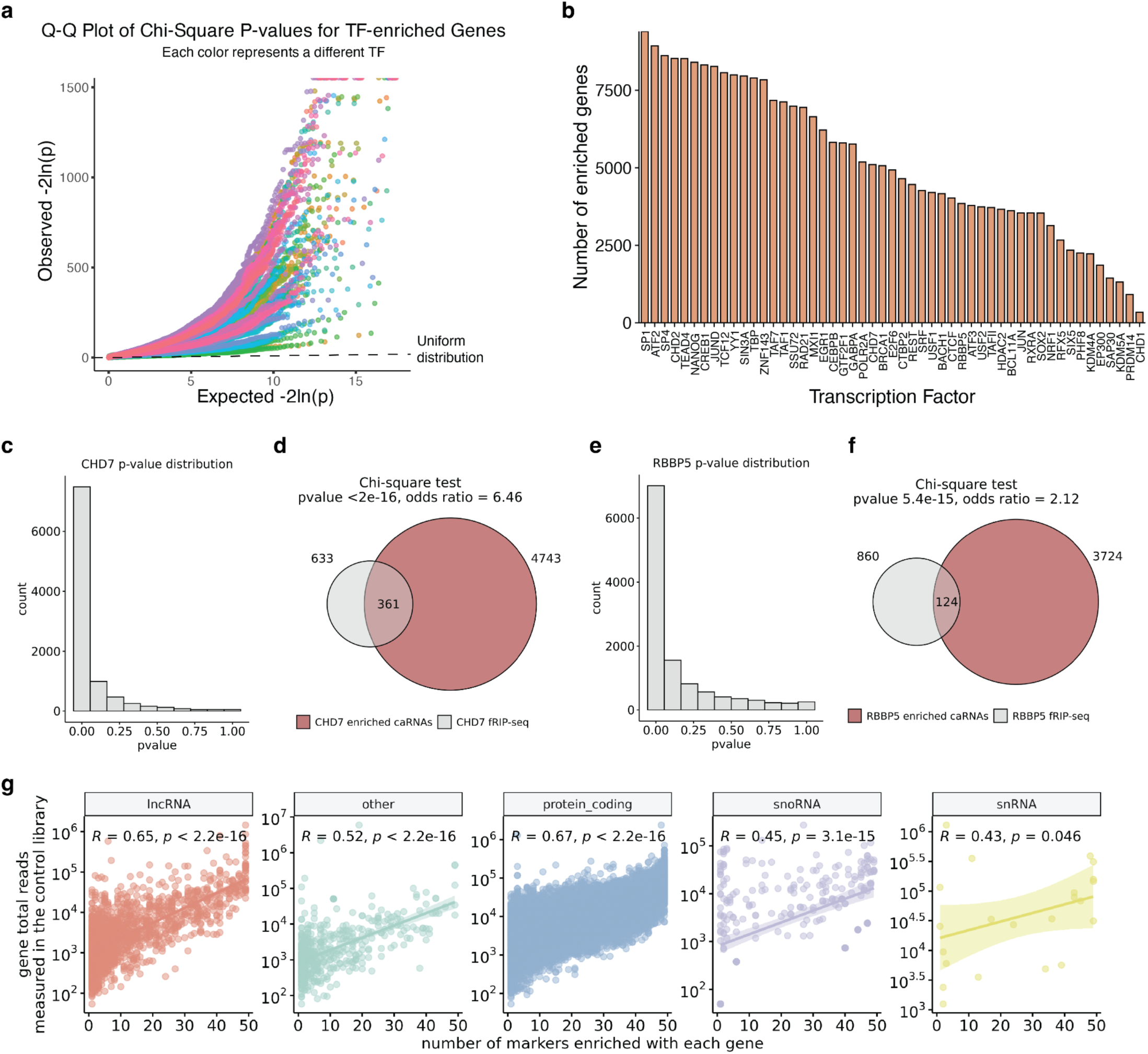
a) Q-Q plot showing the p value distributions of all 52 TF markers (color dots, each color corresponds to one transcription factor) and comparing them to the uniform distribution (dashed line). b) Number of enriched genes for each transcription factor. c) Histogram of p-value distribution from association test. d) Overlap between CHD7 TFBS enriched caRNA species and CHD7 fRIP-seq identified CHD7 protein targeting RNAs. e-f) Similar to c-d) but for RBBP5. g) Relationship between number of caRNA associated transcription factors and gene total transcribed read counts in untreated control library.

**Table S1.** iMARGI libraries overview.

| **Library** | **Total Read pairs**<br>**(de-multiplexed)** | **Total deduplicated mapped reads**<br>**(both ends mapped)** | **MAPQ>=30 and unique**<br>**mapping** |
| --- | --- | --- | --- |
| RNase 1 | 719,424,571 | 409,443,906 | 48,926,803 |
| RNase 2 | 950,806,144 | 344,012,782 | 50,633,451 |
| RNase 3 | 1,519,702,193 | 835,597,061 | 92,517,185 |
| Control 1 | 1,009,524,877 | 783,821,122 | 134,185,427 |
| Control 2 | 297,811,325 | 184,538,199 | 25,330,555 |
| Control 3 | 330,531,513 | 202,604,067 | 16,224,875 |
| Control 4 | 1,004,910,451 | 587,454,453 | 42,362,018 |
| Control 5 | 922,845,725 | 726,916,769 | 123,383,403 |

